# Regulation of the Molecular Shuttle Service: A Multiscale Model of Aquaporin 2 Recycling

**DOI:** 10.1101/2021.06.22.449377

**Authors:** Christoph Leberecht, Michael Schroeder, Dirk Labudde

**Affiliations:** University of Applied Sciences Mittweida, Mittweida, 09648, Germany; Biotechnology Center (BIOTEC), TU Dresden, Dresden, 01307, Germany

## Abstract

The response of cells to their environment is driven by a variety of proteins and messenger molecules. In eukaryotes their distribution, which is regulated by a vesicular transport system, is important for a tight cellular response. The recycling of aquaporin 2 between membrane and storage region is a crucial part of the body water homeostasis and its malfunction can lead to Diabetes insipidus. To understand the regulation of this system, we aggregated pathways and mechanisms from literature and derived three models in a hypothesis-driven approach. Furthermore, we combined the models to a single system to gain insight into key regulatory mechanisms. To achieve this we developed a multiscale computational framework for the modeling and simulation of cellular systems. The analysis of the system rationalises that the compartmentalization of cAMP in renal principal cells is a result of the protein kinase A signalosome and can only occur if specific cellular components are observed in conjunction. Endocytotic and exocytotic processes are inherently connected and can be regulated by the same protein kinase A signal.

## Introduction

The response of eukaryotic cells to signaling molecules in adjacent tissues is a complex interplay of many molecular systems^1^. In recent decades, research has made phenomenal progress concerning timing, intensity, and triggers of signaling events. The response is frequently linked to the transport of chemical entities via intracellular vesicles^2^. Vesicles are mobile membrane enclosed compartments that are able to move along filaments of the cytoskeleton^3, 4^. The surface of a vesicle is a mosaic of proteins that is responsible to attach vesicles to the correct filament, trigger their departure as a response to changes in the surrounding concentrations, and catalyze reactions. For example: Neurotransmitters are stored in vesicles and released upon action potential arrival^5^ and the absence of insulin triggers the internalization of GLUT4 transporters^6^. New imaging techniques have made it possible to observe cells in unprecedented detail, pinpointing location and concentration of cellular components. It became evident, that especially the location of key molecules plays a major role in specificity and sensitivity to external stimuli^7^. However, the integration of all relevant mechanisms in conjunction with their location in the cell into a complete signaling cascade is an intricate undertaking. The modeling process is further complicated by the different scales of time and space. Systems biology is therefore challenged with the integration of mechanistic details and rules from different studies^8^. A model typically consists of multiple components whose collective behavior allows the verification or rejection of a hypothesis. The choice of which components to integrate in a system is strictly speaking a hypothesis in itself^9^. Hence, the underlying structure of the model should be flexible enough to allow for the exchange of model parts, whenever new knowledge arises, or simply to test different concurrent approaches^10^. The modeling of the vesicular transport system is most often tackled by agent-based approaches^11–13^ whereas other intracellular transport and signaling phenomena are approached via continuous models^14–16^. Signaling cascades that involve vesicles for intracellular transport include phenomena of both areas. To tackle this problem, we devised a hybrid modelling and simulation approach that is able to calculate the behaviour in both worlds in a single simulation.

The ability to encapsulate cellular behaviour of different scales into independent modules allowed us to integrate and a large number of mechanisms and parameters. Using data from more than 100 published sources we devised the most complete model of the vesicular Aquaporin 2 (AQP2) transport system to our knowledge.

### Aquaproin 2 recycling

The water balance of the mammalian body is regulated by the interplay of hypothalamus, neurohypophysis, and kidneys^17^. Collecting duct principal cells in the kidney express hormone receptors on their basolateral membranes, that are able to initiate a signaling cascade upon vasopressin binding^18–20^. This results in an increased water absorption at the opposing apical cell membrane^21^. One of the main components of this cascade is the water transporter AQP2^22, 23^. In its basal state, a large portion of the transporter is stored in vesicles in the perinuclear region of the cell^4^. A phosphorylation of AQP2 by cAMP-activated Protein kinase A (PKA) results in the attachment of the vesicle to actin filaments and a subsequent transport to the apical cell membrane^24, 25^. Concurrently, AQP2 is reclaimed via clathin-mediated endocytosis^26^, resulting in a dynamic distribution of the water-transporting protein^27, 28^. A detailed review of the involved mechanisms can be found in Supplementary Information 1.

### Motivation and Aim

We investigated biological phenomena that are involved in the vesicular recycling in general and in the AQP2 response especially:

- The unique nature of PKA regulation^29^
- Organization of proteins in signalosomes^30^ and biological condenstates^31^
- cAMP compartmentalization and its apparent low diffusivity^32^
- Coupling of microscopic and macroscopic signal transduction^33^

We found that all these different aspects are inherently tied together and are key elements to understand the PKA/AQP2 signalosome. PKA is extensively being investigated for over 50 years, and still new aspects of its mechanisms are unveiled almost yearly. Recently, it was uncovered that the autophosphorylation of Protein kinase A regulatory subunit (PKAR), happens in the absence of cAMP^34^ and intricate allosteric processes release Protein kinase A catalytic subunit (PKAC) from PKAR^35^ upon cAMP binding. Furthermore, anchoring proteins such as A-kinase anchoring protein (AKAP) are important for the localization of protein kinases. In renal principal cells AKAP 18*δ* is associated with AQP2 bearing vesicles and provides binding sites for phosphodiesterase type 4D3 (PDE4)^36^ and Serine/threonine-protein phosphatase 2B (PP2B)^37^, which are proteins required for signal modulation and termination. Weak and strong binding interactions^38, 39^ between the components of the so called PKA signalosome are deemed increasingly important for the signaling process by increasing local cohesion of the proteins^30^. In recent years, it was discovered that many proteins are able to from biomolecular condensates^31, 40^; a complex spatial and temporal interplay of molecular components^30^. Molecules that are part of condensates are exposed to drastically different environmental conditions compared to globular proteins in solution.

The apparent diffusivity of the small signaling molecule cAMP was observed to be magnitudes slower than molecules of comparable size, when observed in vivo^41^. Furthermore, there are areas in the cell where cAMP is present in concentration gradients that should intuitively not be possible considering the physical properties of the cytoplasm. This so called compartmentalization of cAMP is crucial for the activation of PKA^42–44^. Interestingly, most studies found that those requirements are very restrictive^43, 45^. The processes required for AQP2 migration to the cell membrane rely on cAMP compartmentalization^46^. We propose that PKA and multiple associated proteins form condensates that regulate cAMP influx and the subsequent activation cascade for PKA. To consider this hypothesis, we implemented models and tested the conditions and components required for the compartmentalization and activation of the AQP2 signaling pathway. We reviewed literature knowledge of this signaling cascade and found qualitative as well as quantitative building blocks. This allowed us to verify applicability and parameters of the individual components. Furthermore, we were able to deduce cellular behavior in parts of the model where no information could be determined from literature. This knowledge allows to suggest new experiments where research can focus to fill in gaps and approach whole cell models in an incremental approach^47^.

There are excellent toolkits available for the modeling of reaction-diffusion systems^48, 49^ as well as for the agent- or algorithm-based nature of macroscopic cellular behavior^50^. In order to address systems that involve both macroscopic changes, such as the movement of vesicles, as well as microscopic changes resulting from chemical processes a hybrid model is required. We therefore developed an integrated approach that allows for hypothesis driven modeling of complex signaling pathways. Furthermore, we implemented this approach in a framework that encourages the definition of a rule based reaction- and behavioral systems. We build the models sequentially, starting with the PKA signaling response. Afterwards the model was extended to include cAMP diffusion and compartmentalization. Finally, endo- and exocytosis, as well as intracellular transport were added. After each step, we took stock and evaluated the behavior of each model individually and in conjunction with the previous results.

## Results

### Allosteric phosphorylation model

The group of Susan Taylor^35, 51, 52^ systematically explored the mechanisms of PKA activation. New models are already starting to include and explore this mechanistic knowledge^34^. We implemented the most recent model for the activation of PKA and included the subsequent phosphorylation of AQP2 and PDE4 (see Fig. 2). For a detailed description and biological background of the model see the Supplementary Information 1 Section 1. The most interesting observation in the recent years is, that the active subunit of PKA, PKAC, binds to the regulatory subunit PKAR with a high affinity and phosphorylates it. In principal cells the regulatory subunit that regulated PKA response at vesicles is PKAR II*β*^53^. The phosphorylation will prevent binding of further PKA, but the PKA that phosphorylated the PKAR is trapped in the bound state until two molecules of cAMP bind to PKAR. This leads to an interesting dynamic, where the PKAC/PKAR complex acts like a loaded spring that is able to release PKAC if enough cAMP is available and needs to be compressed again to initiate the subsequent response. We explored the reaction parameter space of this phosphorylation model, by observing the phosphorylation ratio of AQP2 and the PKA activity in 5184 different settings for five minutes each.

**Figure 1.**
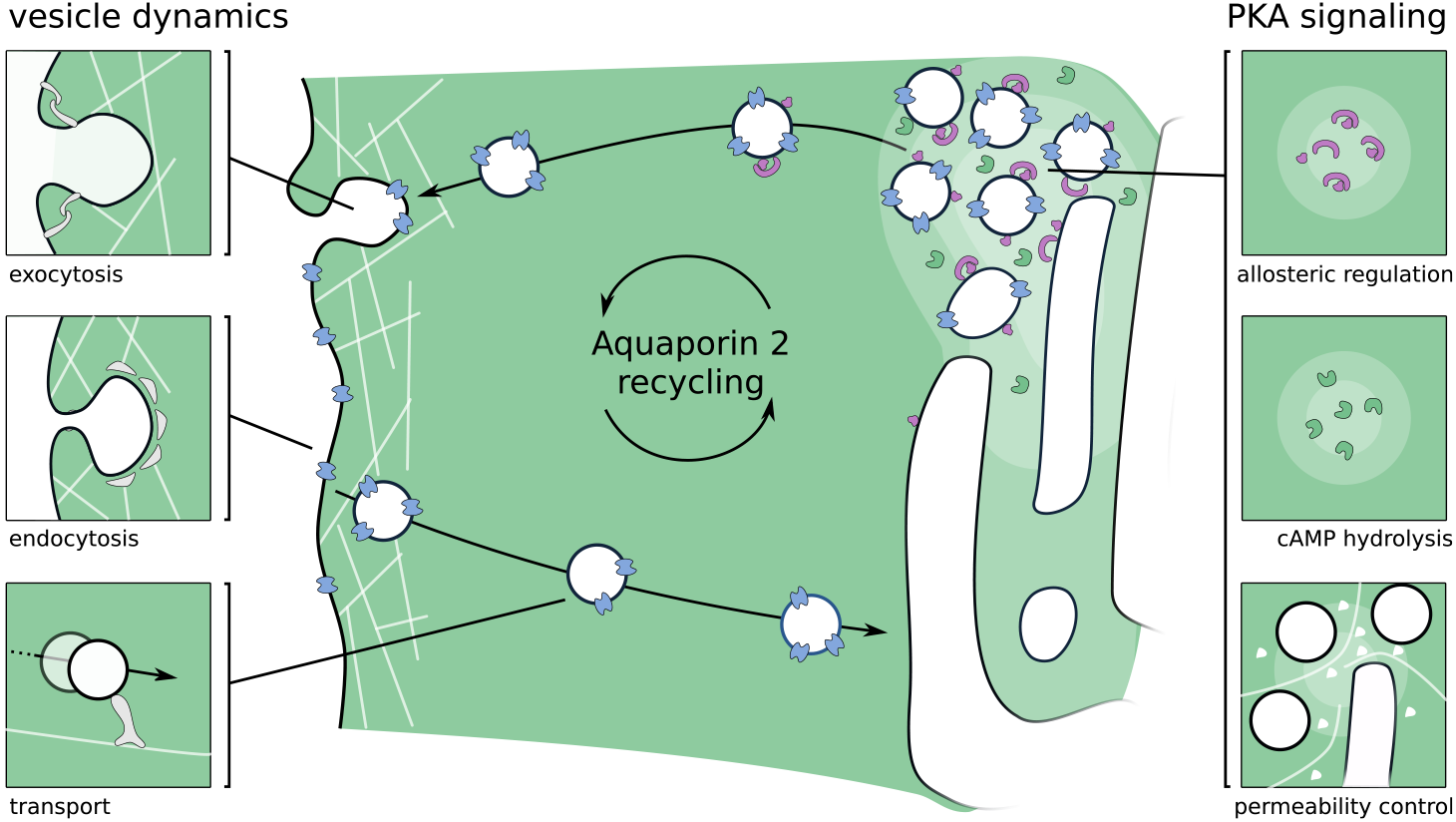
Graphical abstract: Key mechanisms involved in AQP2 recycling. Macroscopic effects are displayed on the left side of the figure, whereas microscopic effects are displayed on the right. AQP2 positive vesicles are transported from the intracellular storage region to the apical membrane upon cAMP signal.

**Figure 2.**
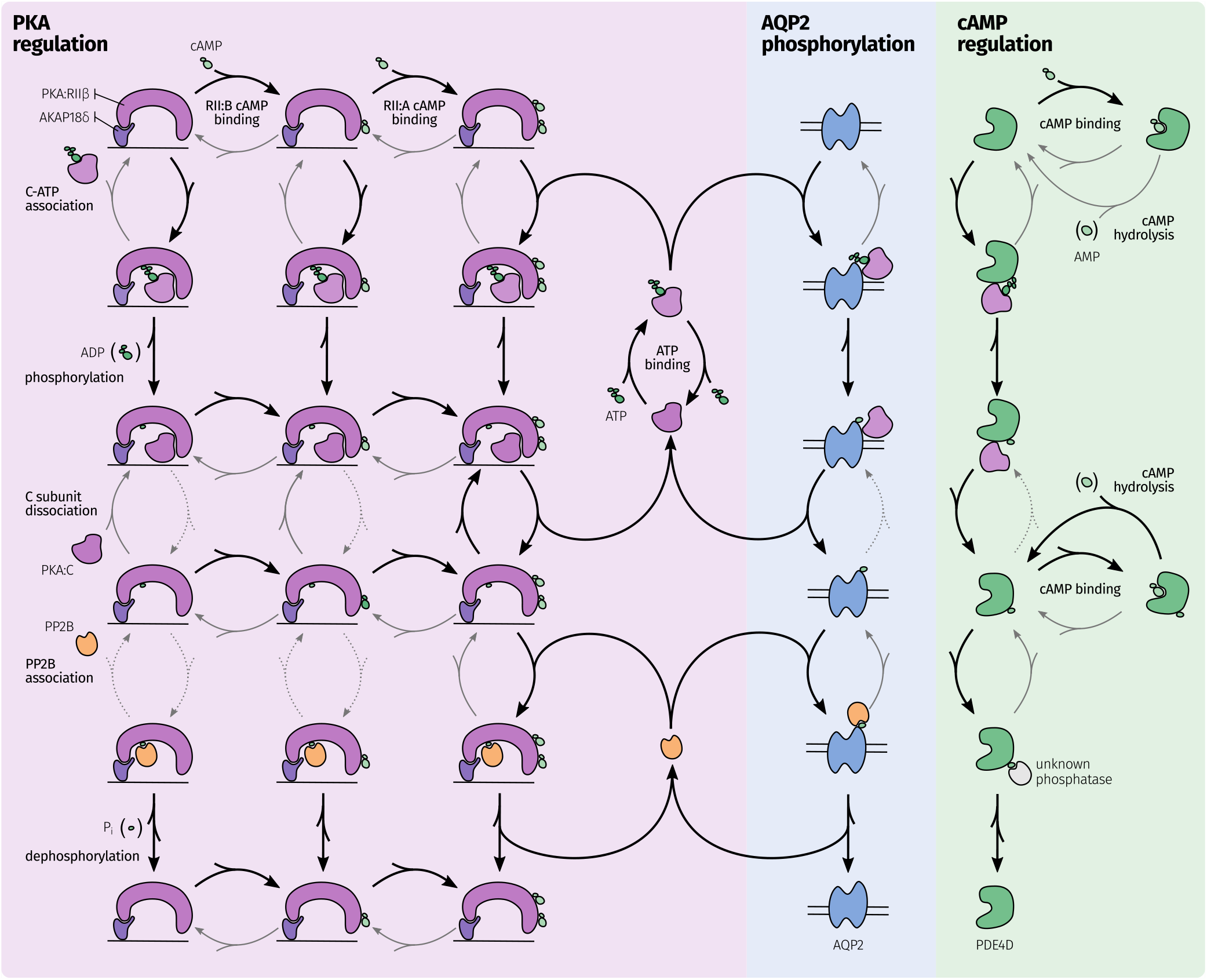
PKA regulation and effect on AQP2 and PDE4 phosphorylation. The network of possible reactions during signal processing as a response to cAMP. Black arrows indicate preferred and fast reactions, gray arrows show slow but still significant reactions, gray and dotted arrows are considered negligible in their frequency. The following reactions in each row from top to bottom: PKA and substrate association, substrate phosphorylation, substrate dissociation, phosphatase and substrate association, and PP2B phosphorylation and release. Substrates are only displayed once and considered implicitly row and column wise. PKAC is only released upon binding of a second cAMP to the regulatory subunit. A negative feedback loop is present, where the released PKAC phosphorylates PDE4, leading to an increased PDE4 activity which decreases the cAMP concentration.

#### PKA activation is buffered and underlies a positive feedback loop

The simulations show, that PKA activity is highly sensitive to the influence of cAMP (see Fig 3B). cAMP causes PKAC to be released from its regulatory subunit, and subsequently free PKAC is able to phosphorylate PKAR, reducing its affinity to PKAC^51^. This positive feedback loop increases the amount of unbound PKAC. Since PKAR is available in excess^38^ and since two molecules cAMP are required to release PKAC, PKAR is able to buffer some cAMP and the amount of unleashed PKAC is initially small. Nevertheless, the buffering of cAMP is not able to significantly alter the process of PKA activation once enough cAMP is available. The process might be required to attenuate volatile cAMP concentration in the cell to inhibit premature activation of the positive feedback loop.

**Figure 3.**
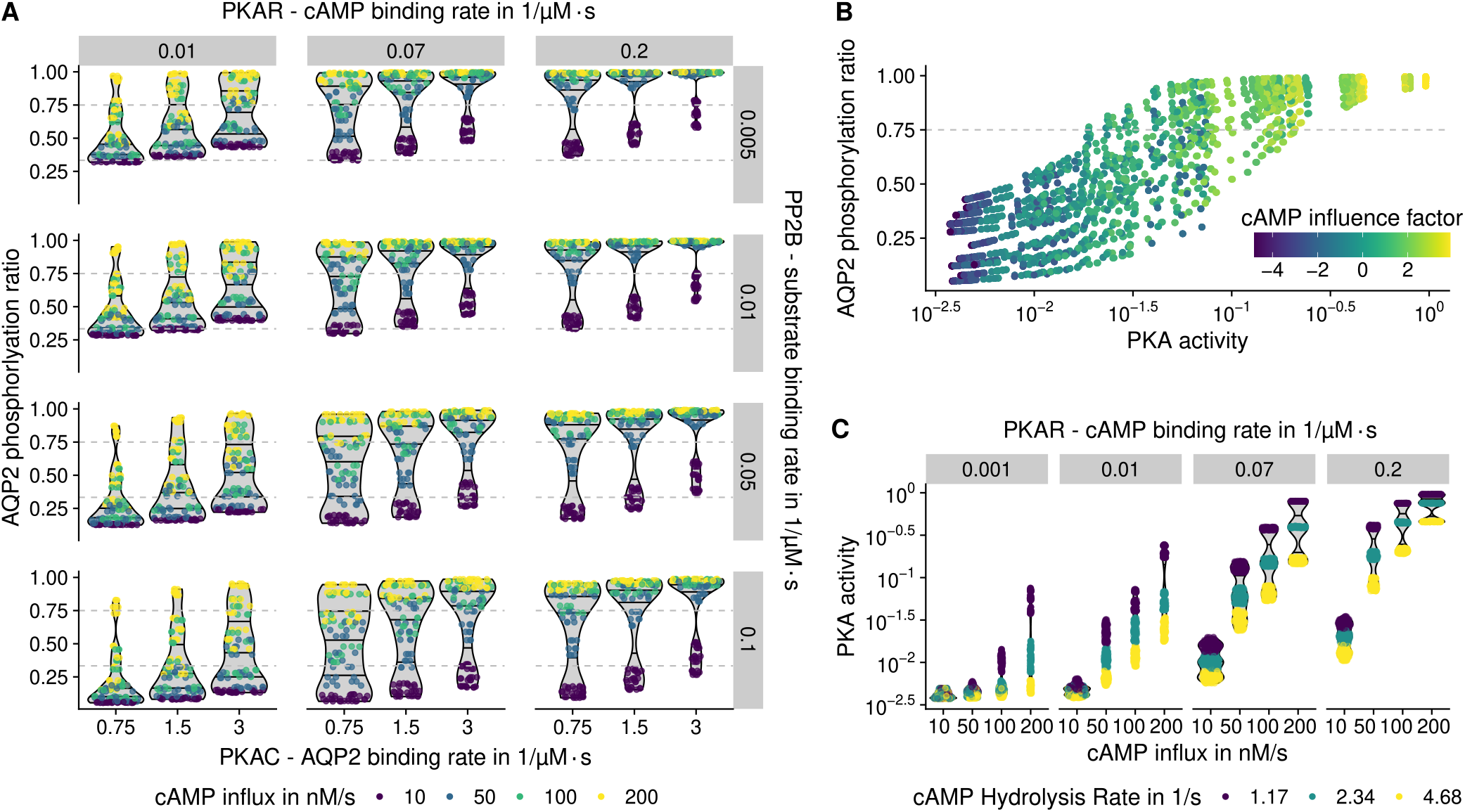
Phosphorylation of PKA and AQP2. Each point indicates the corresponding value after 5 minutes simulation with varying parameters. Basal phosphorylation and threshold AQP2 phosphorylation ratios are shown as gray dashed lines. **A**: High cAMP influx is a major determinant for phosphorylation. Depending on PP2B binding rate (negative effect) and PKAR binding rate (positive effect) the response is shifted. **B**: cAMP influence factor is calculated by log(cAMP influx · cAMP binding). High cAMP influence on the simulation leads to high PKA activation and subsequent AQP2 phosphorylation. **C**: cAMP hydrolysis by PDE4 significantly influences PKA phosphorylation, depending on the cAMP influx and PKAR - cAMP binding rate.

#### Hydrolysis of cAMP decreases PKA activation

Phosphorylation of PDE4 has the possibility to initiate a negative feedback loop by increasing cAMP hydrolysis^54^, subsequently reducing the PKA activation level. In this model and with parameters varying around physiological conditions a negative feedback loop could be observed. PDE4 activation and hydrolysis rate had a big impact on PKA activity (Fig.3C), but played a secondary role when considering AQP2 phosphorylation. Further, PKA activity is heavily influenced by the cAMP influx and the PKAR binding rate (see Fig. 3B). About 75% of the AQP2 molecules phosphorylated at S256 are sufficient to trigger the departure of the vesicles to the apical membrane^55^. Therefore, we used this threshold to evaluate whether the phosphorylation is sufficient for the propagation of the signal. Using this setup, an influx of 200 nMs^-1^ resulted a sufficient phosphorylation ratio in all simulations where cAMP binding rate was 0.07 μM^-1^ s^-1^ or greater.

#### Phosphatase PP2B reverses PKA response

The binding rates of PKAC and PP2B to AQP2 are the deciding factors for the ratio of AQP2 phosphorylation (see Fig 3A). High binding rate of PKAC and low binding rate of PP2B lead to an increased AQP2 phosphorylation ratio at similar PKA activity levels. PKAC is able to phosphorylate PKAR, PDE4 and AQP2 in this model and additional targets *in vivo*. This competitive binding was considered by using two reactions for binding and phosphorylation, essentially trapping a small amount of PKAC in each time step. Decreasing the binding rate of PKAC to AQP2 therefore increases the amount of PKAC available to phosphorylate other components. The variation of the relevant binding affinities was unable to produce an effective difference in PKAC concentration, and has therefore minor impact on PKA behavior. PP2B is indirectly activated by Ca^2+^ via Calmodulin. Since the Calmodulin pathway was not considered in this model, the parameter variation in PP2B binding rate is used as a proxy to determine the influence of Ca^2+^ on the PKA/AQP2 pathway. High PP2B binding rates were able to reduce AQP2 phosphorylation in some models, but high cAMP influx was able to overcome this effect for nearly all setups (see Fig 3A).

We demonstrate that the model is able to represent multiple aspects of PKA activation. The excess of PKAR and two cAMP binding sites prevent premature activation. An initial positive feedback loop wherein PKAR phosphorylation reduces PKAC binding leads to a switch-like activation of all phosphorylation targets of PKA. The subsequent negative feedback loop involving PDE4 is able to effectively reduce cAMP concentration and PKA activation. The simulations were performed without spatial components that could prevent cAMP from reaching its destination. In the next models we wanted to investigate, how cAMP compartmentalization influences the PKA and AQP2 phosphorylation.

### cAMP compartmentalization in the vesicle storage region

One of the most well known second messengers cAMP is compartmentalized frequently in signaling pathways^42–44^. For this to occur, various chemical and physical prerequisites have to be met^56^. The phenomena that influence the compartmentalization of cAMP can be condensed to three major factors: The hydrolysis of cAMP by phosphodiesterases, the apparent permeability of the cytoplasm (also known as diffusive restriction), and transient or permanent chemical interactions^43, 45, 57, 58^. We modeled and assessed these factors by a variation of kinetic parameters and environmental setup. The permeability of the cytoplasm was explored by introducing macroscopic objects that block access to the areas where PKA is present and additionally by defining a permeability parameter that slows diffusion in those storage regions (see Figure 4). The hydrolysis of cAMP was explicitly modeled and binding affinities to all relevant components was systematically explored. The detailed model setups and parameter variations are listed in the Supplementary Information 1 Section 2. To evaluate the importance of cAMP for the actual regulation of the AQP2 pathway, both, the developing cAMP gradient as well as the PKA activity need to be considered.

**Figure 4.**
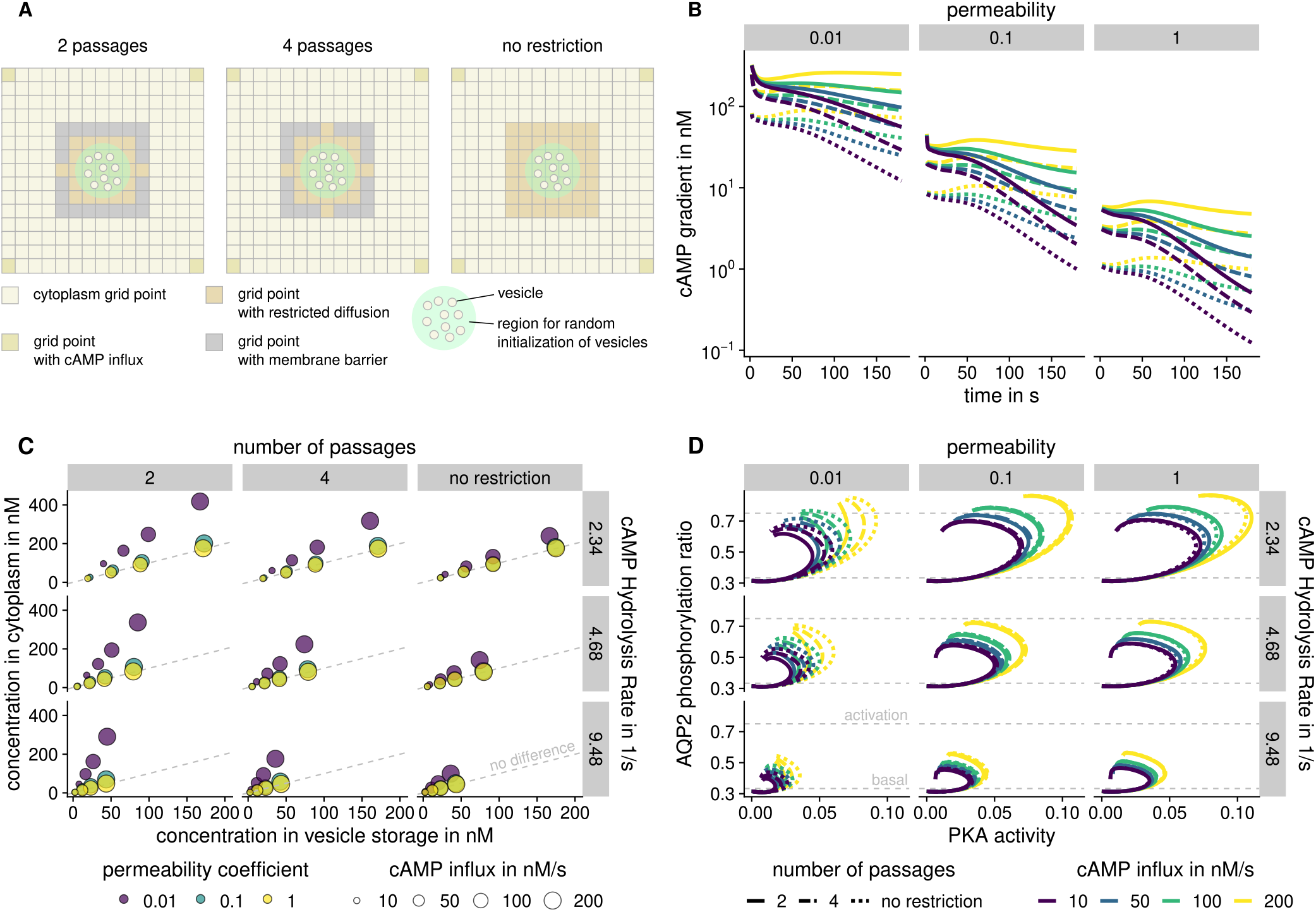
cAMP compartmentalization in different environments. **A**: Schematic representations of restricted environmental setups with and without barriers around a vesicle storage region. **B**: cAMP gradient over time under varying conditions. Simulations with no permeability reduction are unable to maintain effective gradients. **C**: Influence of PDE4 hydrolysis rate and number of barrier passages on concentration in different compartments after five minutes of simulation. Circle size indicates cAMP influx, whereas color shows different permeabilities. Only the combination of diffusive restriction and low permeability was able to create significant differences in concentration. **D** High PDE4 hydrolysis rate leads to lower cAMP concentration in the vesicle region. High cAMP influx allows to overcome the phosphorylation threshold.

#### cAMP compartmentalization only develops in regions with reduced permeability

The most important factor for the emergence of gradients is permeability (see Fig. 4B). It is possible to observe compartmentalization effects in all simulations. Differences on the scale of 10-fold reduction as suggested by Iancu and colleagues^59^ are only obtained with low permeability and high cAMP hydrolysis rates. Simulations without any obstacles and full permeability were unable to attain a significant cAMP gradient, even when considering highly potent phosphodiesterases and buffering (see Fig. 4C). With high permeability, the whole system is affected by the cAMP reduction, resulting in a nearly uniform distribution across compartments. Others have also observed these effects in experiment and simulation in neonatal cardiac myocytes^45, 60^, dendrites of neurons^57, 61^ and other cell types^56, 62^. Stephan and colleagues observed cAMP compartmentalization in renal principal cells^46^. Since cAMP is produced at the basolateral membrane and vesicles are stored close to the apical membrane, cAMP needs to travel through the cytoplasm. It seems feasible that a locally reduced cAMP concentration is required to prevent premature activation of PKA as a response to the basal cAMP concentration^41^. After the cellular concentration of cAMP increases due to activation by vasopressin, the actual response pathway is triggered. The components for this to take effect are all coupled to the AQP2 vesicle^36, 37^.

#### Buffering effects play a minor role in compartmentalization

Buffering was proposed to have effect on cAMP compartmentalization. cAMP buffering is the capacity of proteins and other cAMP binding molecules to temporarily or permanently fixate cAMP rendering it unavailable in the pool of free cAMP. In this setup buffering is mostly performed by PKAR subunits. Since PKAR is available in excess, it is able to bind a significant amount of cAMP (twice the concentration of PKAR in the system)^38^. PDE4 buffers a negligible amount just before catalysis. As discussed in the phosphorylation model, buffering is able to create short term reduction of the cAMP and therefore fine tune and stabilize small fluctuations in the sink. However, binding to PKAR is also required to activate PKAC. Therefore, cAMP buffering at PKAR can not be viewed as a means to generate cAMP gradients that result in a specialized response. PKAR and PDE4 are only a part of the possible binding partners of cAMP. It would be interesting to analyze the specific and unspecific binding of cAMP to gain more insight as to how unspecific buffering contributes to compartmentalization.

#### The creation of cAMP sinks is a delicate balance between multiple factors

PDE4 degrades cAMP in the cell. PDE4 in the PKA signalosome can be activated by PKA, which increases its cAMP hydrolysis rate^54^. A high hydrolysis rate leads to lower cAMP concentrations close to the vesicles. The creation of cAMP sinks is a delicate balance between the amount of cAMP that is produced, the turnover rate of PDE, and the reduced access of cAMP to relevant regions of the cell (see Figure 4C). Physiologically, increasing the cAMP influx seems inefficient, since large amounts of energy would be required to produce cAMP, only to degrade it moments later. The amount of PDE4 required to decrease cAMP influx without diffusive restriction would far exceed physiological ranges^45^. This can be confirmed by our simulations. The cAMP influx is not sufficient to compete with the degradation by PDE4 for permeabilities of 0.1 or higher, leading to a globally reduced cAMP concentration and no compartmentalization. A high diffusive restriction in the vesicle area is an elegant solution to compromise both factors. Fewer cAMP molecules reach the vesicles and whenever the influx exceeds the degradation capacity of PDE, it results in an activation of PKA. Other factors gain influence, if permeability is 0.1 or less. We used the phosphorylation ratio of AQP2 to determine under which conditions cAMP compartmentalization is able to influence signaling. We found that phosphorylation levels of AQP2 surpass the 75% threshold, if the cAMP hydrolysis rate is low enough and cAMP influx is sufficiently high (see Figure 4D).

In conclusion, the prime factor to create effective cAMP compartmentalization was the permeability of the vesicular storage. Hydrolysis of cAMP by PDE4 is able to fine tune the concentration of cAMP and regulate the signaling response. The explicit buffering modeled in this study did not contribute significantly to sustainable compartmentalization.

### Clathrin-mediated endocytosis model

In the basal state of the cell vesicles are located in the storage region until they are transported to the membrane for fusion. New vesicles are created at the apical membrane via clathrin-mediated endocytosis depending on SRC phosphorylation^26, 63^. A constantly shifting imbalance of exocytosis and endocytosis is the major driver behind the water acquisition of principal cells in the kidney. An increased endocytosis shifts the majority of AQP2 to the storage region, whereas increased exocytosis leads to high AQP2 concentrations in the apical membrane. The increase in exocytosis is mediated by the pathways we explored in our previous models. The frequency and regulation of endocytosis is the focal point of the endocytosis model.

#### Endocytosis is mediated by PKA activation and AQP2 concentration

Endocytotic pits, the precursors of clathrin-coated vesicles, emerge spontaneously on the apical membrane surface and the maturation from pit to vesicle is correlated to key cargo molecules^64^. We determined the rate of endocytotic pit formation as well as the cargo collection speed, from the approximate number of AQP2 molecules per vesicle^65^ and parameter variation. Furthermore, we implemented a regulation mechanism involving Non-receptor tyrosine kinase (SRC) to model the AQP2 accumulation observed in active cells^63^. The model to evaluate the parameters for this process imitates a apical membrane section with a surface of one μm^2^ that contains the expected molecules after cAMP stimulation. Knowledge from the phosphorylation and compartmentalization models was used to constrain the parameters of the endocytosis model. The detailed model setups and parameter variations can be reviewed in the Supplementary Information 1 Section 3.

#### Pit emergence and cargo addition are coupled parameters

A low cargo addition rate results in a high number of abortive pits, since the threshold for successful maturation cannot be reached in time (see Fig 5A). Furthermore, high pit formation rates lead to a high number of abortive pits (see Fig 5B). There is an upper threshold to the number of productive vesicles that are able to form, imposed by the total number of AQP2 in the membrane. After a significant amount of cargo has been transferred to vesicles no more productive pits are able to develop. If many pits form in parallel, the available cargo molecules are distributed across all pits in the membrane. Hence, not enough cargo can be accumulated in each pit and the number of productive pits remains low.

**Figure 5.**
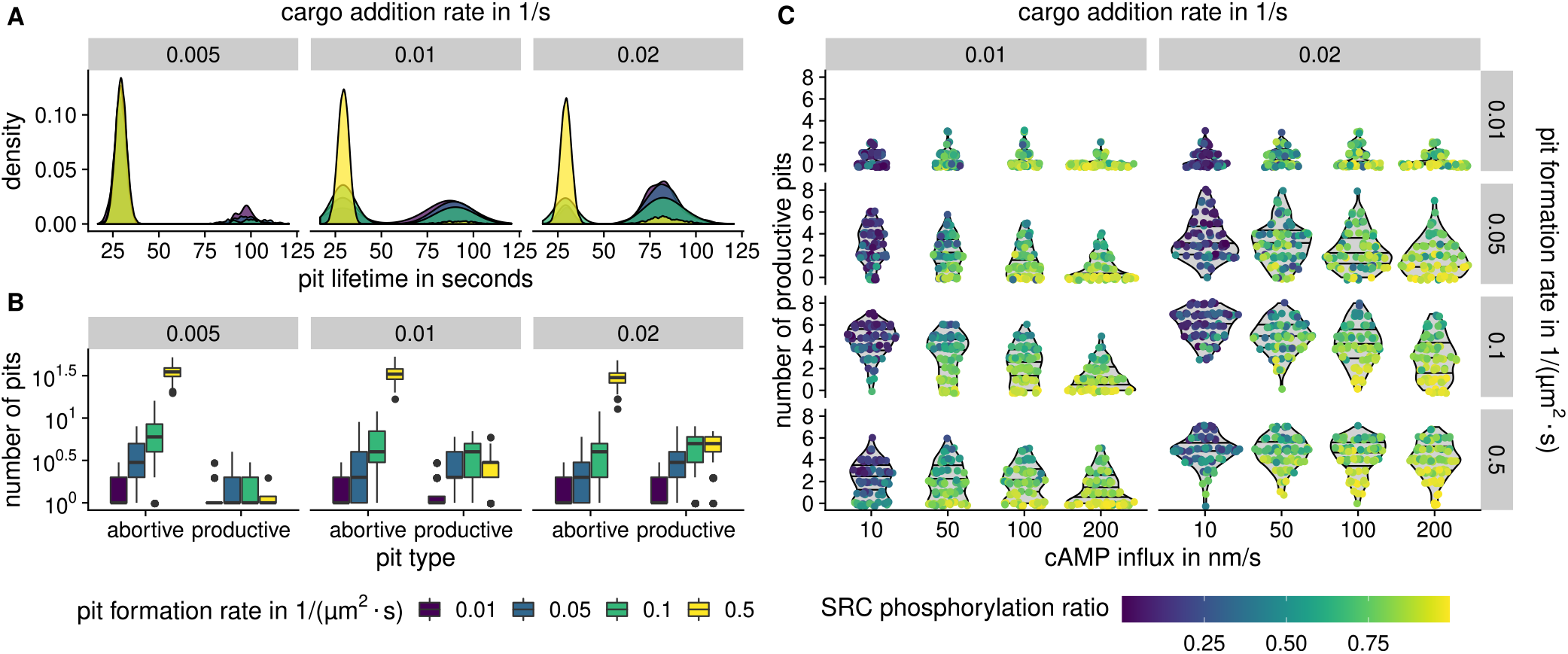
Distribution of abortive and productive endocytotic pits. **A**: Lifetime of endocytotic pits across all simulations at varying cargo addition rates. The first peak indicates abortive pits that did not enter maturation phase, second peak indicates pits that matured to vesicles. **B**: The number of abortive and productive pits after five minutes. High cargo addition rate leads to high number of productive pits. The total number of AQP2 molecules in the membrane exerts an upper threshold for productive pits. **C**: Number of productive pits at varying parameters, colored by average SRC phosphorlyation ratio. Low cAMP influx leads to higher number of productive pits, given a sufficient pit formation rate. High SRC phosphorylation ratio results in fewer productive pits.

#### Model of SRC inhibition is able to reduce productive pit count

In this model the ratio of inhibited and active SRC was used to scale the cargo accumulation rate, such that inhibited SRC leads to an increase in abortive pits. The inhibition of SRC is mediated by a phosphorylation of Tyrosine 527, which is performed by activated C-terminal SRC kinase (CSK)^66^. CSK itself can be activated by PKA via phosphorylation of Serine 364^67^. Therefore, an activation of PKA leads to an inactivation of SRC and subsequently to AQP2 retention in the apical membrane. The influence of SRC phosphorylation can be seen in Fig. 5C. The decrease in effective cargo collection rate resulting from SRC phosphorylation is able to qualitatively reproduce the observations made by Cheung et al.^63^. A high cAMP influx rate is associated with fewer productive pits. Inversely, a high number of productive pits can be observed in systems with low cAMP influx, as a result of inactive PKA and dephosphorylated SRC.

### Full recycling model

For the recycling model we use the knowledge gained in previous sub-models to set up a spatio-temporal model of the vesicular recycling system of renal principal cells. The model represents a subsection of the cell that includes a vesicular storage region as well as the apical cell membrane. The processes and reactions implemented in this model are visualized in Supplementary Figure 1 and detailed in Supplementary Information 1 Section 4.

As our previous models have shown, diffusion restricted regions are required to achieve cAMP compartmentalization. Therefore, two regions are defined that reduce the diffusivity of cAMP (see Figure 6A). Agent based modeling is used to simulate the dynamic behaviour of vesicles. Two exemplary setups are used to evaluate the behaviour shown by the model: activation and recycling. The first activation model exhibits behaviour close to that of natural principal cells. In the recycling model SRC and CSK dephosphorylation have been increased, which allows to observe exocytosis as well as endocytosis in action.

**Figure 6.**
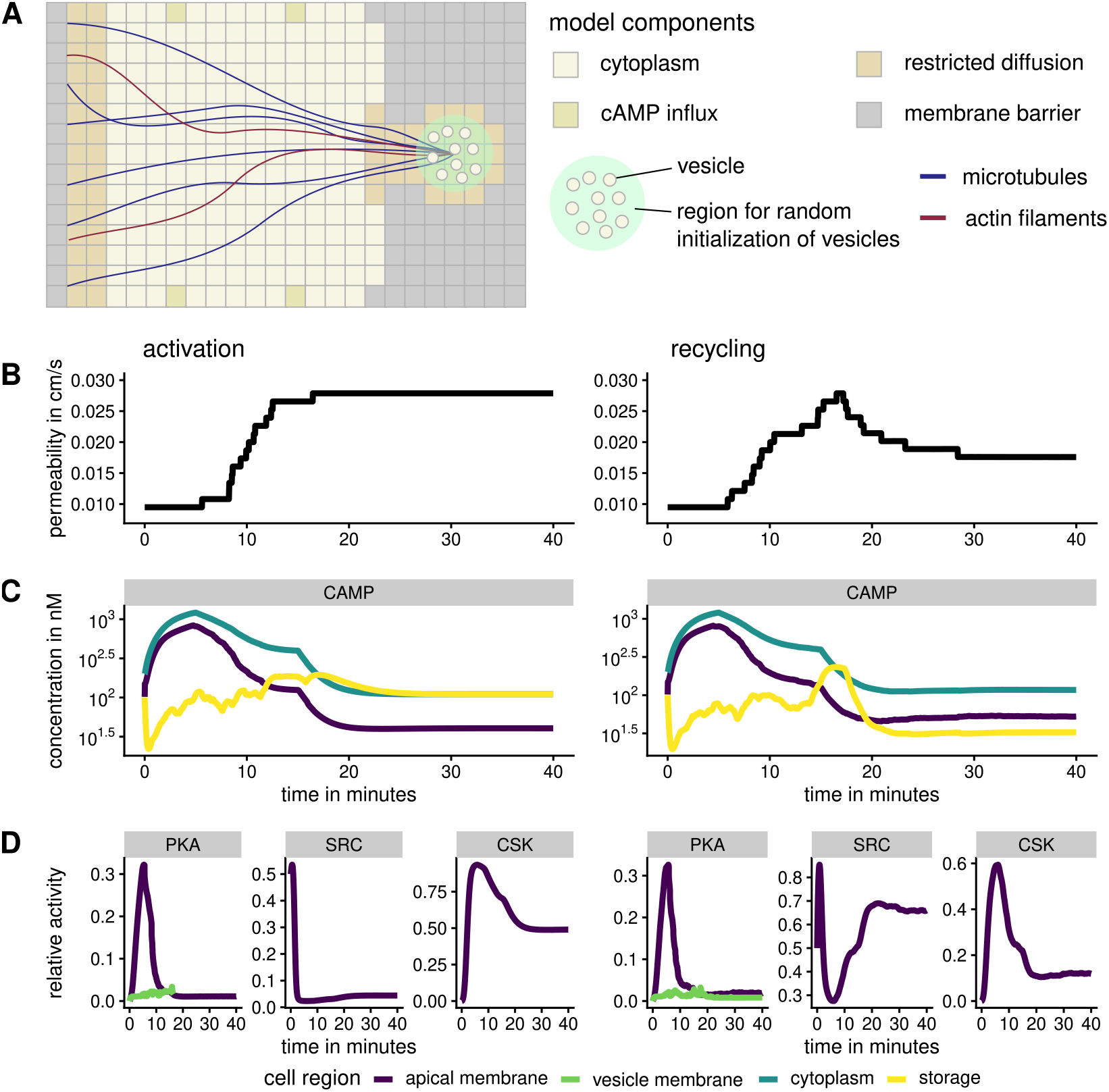
AQP2 transport activation and recycling. Two setups of the full model showcase activation (left) in opposition to recycling (right) when using accelerated SRC and CSK dephosphorylation. **A**: Schematic representation of the environmental setup. The storage region on the right contains the AQP2-positive vesicles that are transported to the apical membrane during simulation. **B**: Membrane permeability estimated from AQP2 concentration in the apical membrane. **C**: cAMP concentration in different cell regions. **D**: Relative activity levels of selected phosphorylation targets.

Both models exhibit a distinct increase in exocytosis. Vesicles are able to move along actin filaments to the apical region of the cell whenever the phosphorylation threshold is reached. Once the vesicle is in close proximity to the apical membrane fusion commences^2^. The initiation of this cascade is regulated by PKA, as demonstrated in previous results. The discrete increases in permeability (see Figure 6B) are the result of individual vesicle fusion events. cAMP produced by adenylate cyclases is simulated by cAMP influx at nodes indicated in Figure 6A. In both models cAMP is created at 400 nMs^-1^ for each of the four cAMP influx grid points for the first 5 minutes and at 200 nMs^-1^ up until 15 minutes. For the rest of the simulation a basal cAMP influx of 50 nMs^-1^ is assumed. Initially, cAMP concentration in the cytoplasm and at the apical membrane increase steadily (see Figure 6C) and the amount of active PKA rises in conjunction. cAMP in the storage region fluctuates significantly, driven by movement and departure of vesicles. After an initial decrease where cAMP binds to PKAR and PDE4 in large amounts, the average cAMP concentration increases slowly. In this model the access to the vesicular storage region is decreased by membranes, whereas the apical membrane is more exposed. The resulting effect is an effective decreased cAMP hydrolysis whenever a major part of the PDE4 concentration is in the storage region. Less cAMP is able to reach PDE4 and fewer cAMP production is required to keep basal cAMP levels stable. Vice versa, the exposure of PDE4 to cAMP is increased whenever vesicles at the apical membrane, decreasing the cellular cAMP stores faster. This mechanism increases the effectiveness of the negative PDE4 feedback loop already present in the initial phosphorylation model and was not explicitly implemented in the model. This is further underlined by the quick deactivation of PKA in the apical membrane, even with significantly increased cAMP concentration (see Figure 6D).

In the recycling model PKA and SRC phosphorylation rise in a similar manner as in the activation model. The increased CSK and SRC dephosphorylation lead to a quick reversal of SRC activity. The increased concentration of AQP2 in the membrane as well as the reactivation of SRC lead to an increased endocytosis rate. Vesicles are then transported to the storage region using microtubule based transport^4^. In the activation model with slower dephosphorylation rates endocytotic events only produce aborted pits and the concentration of AQP2 in the membrane stays constant.

#### Increase in exocytosis and decrease in endocytosis are entwined

cAMP influences not only the exocytosis of AQP2 positive vesicles but also their endocytosis. In the basal state of the cell the majority of AQP2 is kept in storage compartments in the center of the cell. A two pronged approach ensures that the effort of transportation is effectively utilized. The maturation of endocytotic pits is reliant on the cargo concentration in endocytotic pits, hence an increase in AQP2 in the membrane leads to an increase in endocytotic events. The regulation mechanism in place is modeled by SRC inhibition. The actual quantitative influence of SRC to inhibit pit maturation is speculative. It would be interesting to investigate the actual kinetic influence SRC has on pit maturation and/or vesicle scission experimentally. Other mechanisms have been identified that can contribute to the decrease in internalization^68^, therefore it is unlikely that SRC is the only determinant for the inhibition of endocytosis.

#### Molecular condensates can explain localized responses

The diffusive restriction of cAMP and coherence of all molecular components are requirements for a regulated and distinct signal response. The diffusive reduction falls within a regulated margin: If no reduction is present, cAMP renders PKA always active even at basal cAMP levels. Since endocytosis is also reliant on indirect cAMP based activation of SRC kinase, the system transitions to a state where AQP2 concentration in the apical cell membrane is always high. On the other hand systems with high restriction are slow to respond to signals, since few cAMP molecules are able to reach the cAMP storage region at a time, delaying PKA activation. Furthermore, AQP2 does not remain in the membrane, since endocytotic events occur frequently. The decrease in diffusive restriction would therefore be associated with chronically increased membrane permeability, whereas increased restriction would lead to polyuria. All components associated with the AQP2 response are associated in a PKA signalosome. If any of the components would remain in the storage region or at the apical membrane, they accumulate and alter the signal response over time.

#### Deviations between model and phenotype

The original response of principal cells is mostly measured by the water permeability of the apical membrane^23, 69^. Therefore we converted the concentration of AQP2 to permeability as described in Supplementary Information 2. The majority of the activation happens in the first minutes. This is consistent with the previously observed mode of PKA activation, where a positive feedback loop is present. The measurements of Deen and colleagues^69^ show that permeability doubled after 10 minutes and tripled after about 30 minutes when compared to the basal rate. We also observe these ratios, but the measured progression in experiments is closer to a linear gradient. This derivation may have multiple potential origins, of which we will address three that seem the most probable to us. The model describes one storage area and subsection of the cell. Measurements in experimental data were not taken from a single cell but from a cell culture. Potentially, multiple storage sites that are triggered at different times can contribute to a more evenly distributed permeability increase. Furthermore, we are starting the simulation with vesicles that contain a uniform number of molecules. It is probable that the vesicular response is altered depending whether the vesicle was recently recycled or in storage for some time^70^. Additionally, we did not model all the aspects that play a role in the vesicular trafficking, for example the role of Ca^2+^ and the other phosphorylations distinct from S256 have influence on the recycling and alter the specific response^71^.

## Discussion

We combined differential equations and agent-based modeling to gain insight into the vesicular recycling system of renal principal cells. Both approaches compliment each other by modeling different aspects of the vesicular system. Especially the combination of microscopic and macroscopic aspects of clathrin-mediated endocytosis requires the combination of chemical reactions and agent-based modules. We extracted parameters and mechanisms from more than 100 publications and integrated them in one context. Additionally, it was required to work out a viable model of clathrin-mediated endocytosis, extend existing models of PKA regulation, and estimate parameters to link the involved processes.

We categorized the major findings resulting from the models in the following three groups: support for previous findings, new insights, and topics that remain to be explored.

### Support for previous findings

Our model confirms that PKA activity can be tightly regulated by a controlled release and reduced the reassociation of PKAC to PKAR controlled by cAMP as suggested by Zhang and colleagues^51^, and not by phosphorylation of PKAR. We verified that PKA needs to be tethered to the vesicular membrane and to the other components of the signalosome for it to have an distinct effect^38^. Components which are not bound to the vesicular storage region diffuse into the cell and render the response unspecific. The PKA signalosome plays a role in multiple cellular processes, depending on its composition^30^. In renal principal cells at least AKAP18, PDE4, PKAR, PKAC, and PP2B are essential for the cellular response and the formation of condensates seems an important factor in reducing permeability of cAMP.

The phosphorylation of PDE4 initiates a negative feedback loop^72^ in order to resore basal PKA activity. It is being debated if the pathway via cAMP and Ca^2+73, 74^ can be triggered independently and result in comparable AQP2 distributions^63, 75, 76^. We argue that there must be some degree of overlap between the responses since PP2B is also regulated by Ca^2+^ indirectly and that they modulate the reaction in different complementary ways. The fact that both secondary messengers are are connected is known^77^. Nevertheless, two largely uncoupled pathways allow for the regulation on different time scales and/or use cases.

We confirm, that PDE4 concentrations or affinities would need to be beyond physiological ranges^45, 62^ also when viewed in conjunction with the PKA pathway in order to be effective for the generation of CAMP gradients. It turned out that cytoplasmic permeability and, as an result, the ability of cAMP to reach PKA signalosome, is the most critical component that allows for the buildup of cAMP gradients. Nevertheless, no effect in isolation is able to create CAMP gradients the interplay of all effects is required to regulate this phenomenon. The permeability required to create gradients was nevertheless higher than expected. While some gradient can be observed for all permeabilities, effective compartmentalization (on the scale of 10-fold diffusivity reduction) was only consistently obtained for permeabilities ≤ 0.01. Effects that lead to a diffusive slowdown can be the result of the cytoplasmic matrix, cellular crowding, and weak binding interactions^78–80^. Whether these factors are able to create another order of magnitude difference in effective permeability is debatable. A promising explanation comes in the form of liquid phase separated compartments, also known as biological condensates^31^. Biological condensates have the ability to reduce diffusion^81, 82^ of the involved components. Molecules that experience weak and strong binding effects tend to cluster together and promote phase separation. These binding effects can be observed for the majority of proteins involved in the phosphorylation cascade^30, 36–39^. Our study supports this view, that challenges the textbook model of local degradation of cAMP as the major driver of compartmentalization. Furthermore, it could explain the substantial difference in cAMP required for the activation of PKA *in vivo* vs *in vitro*^41^.

The regulation of either endocytosis or exocytosis has little effect on AQP2 accumulation. Both pathways are self regulating to an extend and only the combination of increased exocytosis and decrease endocytosis is able to initiate a distinct response^83^.

### New insights

The modeling and simulation process unveiled that PKAR excess decreases substrate specificity^84^, simply by reducing the probability of PKAC encountering and engaging with other substrates, which could contribute to the apparent 10-fold reduction of the activation constant^59^. In connection to this we also found, that different affinities for PKA phosphorylation targets PKAR, PDE4 and AQP2 had no significant impact on the signal response. Using high PP2B activity as a proxy for Calcium influence indicates that Ca^2+^ does not significantly alter the course of the PKA/AQP2 phosphorylation pathway as well. While the PP2B lowered the basal phosphorlyation rate of AQP2, it was not able to effectively counteract the PKA response. The buffering effects of PKAR are negligible for long term gradient generation and only able to attenuate small fluctuations of cAMP.

All three factors influence the compartmentalization of cAMP slightly differently (see Figure 4C): The cAMP hydrolysis rate mainly affects the cAMP concentration in the vesicle region. The diffusive reduction maintains the cytoplasmic concentration of cAMP, and the cAMP influx impacts both cytoplasm and storage compartments. This allows for a largely independent control of of different spatially distinct cAMP sinks in the cell by using different variants and concentrations of the components that are part of the signalosome. As an result, the combination of components in each cell type is able to determine the specificity and diversity of responses. Furthermore, the apparent efficiency of cAMP hydrolysis is decreased whenever PDE4 is in the storage region. The transport of the PKA signalosome in conjunction with the vesicles exposes PDE4 to the cytoplasm and apical membrane, where PDE4 is exposed to higher concentrations of free cAMP. This increases the effectiveness of the negative PKA feedback loop. This tight regulation of the universal kinase PKA leads to a distinct response, whereas the down regulation can be regulated by more specific components such as CSK and SRC.

The cargo addition rate to an endocytotic pit as well as the emergence of new pits need to be balanced to support formation of productive pits. The cascade of PKA/CSK/SRC phosphorylation proved to be a suitable model to inhibit AQP2 vesicle endocytosis and subsequent accumulation of AQP2 in the apical membrane.

### Unresolved phenomena

The influence of the Calmodulin pathway and the concrete role of Ca^2+^ remain elusive. We speculate, that cAMP is responsible for the activation of the PKA-based response and Ca^2+^ is able to counteract it through PP2B, as well as trigger it independently. The modeling of Ca^2+^ would encompass cAMP producing adenylate cyclases, that are inhibited by Ca^2+85, 86^, as well as Calmodulin^73, 87^, and Myosin^88^. Another pathway involving Short transient receptor potential channel 3^89^ is a promising candidate to explore for apical AQP2 accumulation.

How exactly the apparent low cytoplasmic permeability close the vesicle storage region is maintained is not fully understood. Biological condensates seem to play a major role in creating these phase separation effects. Very recently this phenomenon was found to be critical for PKA regulatory subunit RI*α*^90^. Even if in our setup PKA RII*β* was unsuitable to efficiently regulate cAMP buffering, the different treatment of cAMP diffusion close to vesicles was necessary and lead to distinct PKA activation.

It is becoming more clear that the isolated observation of components *in vitro* can be a bad proxy for their actual behaviour *in vivo*^41^. With the concept of biological condensates in mind, the determination of the individual components rate constants is not enough. It is therefore crucial to determine the influence of “signalosome-partner” proteins to create reliable models. The models lead to the conclusion that the phenomena of cAMP compartmentalization and PKA activation are tightly coupled and need to be viewed in conjunction.

Even though this model includes major aspects of the AQP2 transport pathway, it is by no means complete. During the modeling process we encountered processes that provide substance for further research: Experimental evidence is required for the regulation of vesicle maturation during endocytosis in the context of signaling cascades. How does the cargo concentration influence the vesicle maturation process^91^? How do Src^63^ and Sipa1l1^68^ contribute to the membrane accumulation of AQP2? This and other questions can aid the incremental formulation of a whole cell model of renal principal cells.

## Methods

The modeling and simulation of cellular spatio-temporal systems involves five major aspects:

- The microscopic components,
- the distribution of those components,
- the macroscopic components,
- the location of those components, and
- the definition of their behavior.

Although, the actual components and their behavior differs depending on the system that is to be modeled, the general nature of the components remains the same throughout all cellular systems. We have developed a framework that abstracts the notion of microscopic and macroscopic components and their interactions. This allows the user to focus on modeling the biological system. The following chapters will outline the system setup and background. A detailed description of each aspect can be found in the Supplementary Methods. The simulation and modeling framework SiNGA developed for this work is available at: https://github.com/singa-bio/singa. Version 0.7.0 was used to simulate models presented in this work.

### Definition of microscopic components

We use the term chemical entity to refer to chemically distinct species of molecules (according to Systems Biology Markup Language (SBML) terminology). Chemical entities are structured objects that are the source and the result of binding, release, addition, or removal processes performed by chemical reactions. Chemical entities can be represented by a graph structure, where the nodes are also chemical entities and edges are covalent or non-covalent connections between them. The smallest possible chemical entity is a single node that might represent the regulatory subunit of PKA or the messenger molecule cAMP. Using a rule based definition, chemical entities can be combined to create more complex ones, a process which provides both: the possible reactions as well as their complex reactants.

### Definition of macroscopic components

The simulation space represents a pseudo three-dimensional slice of a biological system with arbitrary width and length, but a fixed height. Initially, the simulation space is compartmentalized into a rectangular mesh to prepare the subsequent application of finite-difference methods. Hereby, the total area of simulation is segmented into a predefined amount of rows and columns. The tiling is further subdivided to compartments by membranes (see Figure 7). A membrane can be placed at the border between two grid points. A single compartment is assigned to each grid point that is not bordered by a membrane, whereas two compartments are assigned to grid points adjacent to a membrane. Hereby, one compartment represents the concentration of entities in the membrane, whereas the other represents the concentration in the remaining space. Each compartment contains a set of chemical entities that are assigned a floating point number based on their concentration in this space. Each compartment is considered well mixed. The space can be setup using raster images (e.g. PNG files), where each pixel represents a grid point and colors represent different compartments. Areas of the same color are automatically surrounded by a membrane.

**Figure 7.**
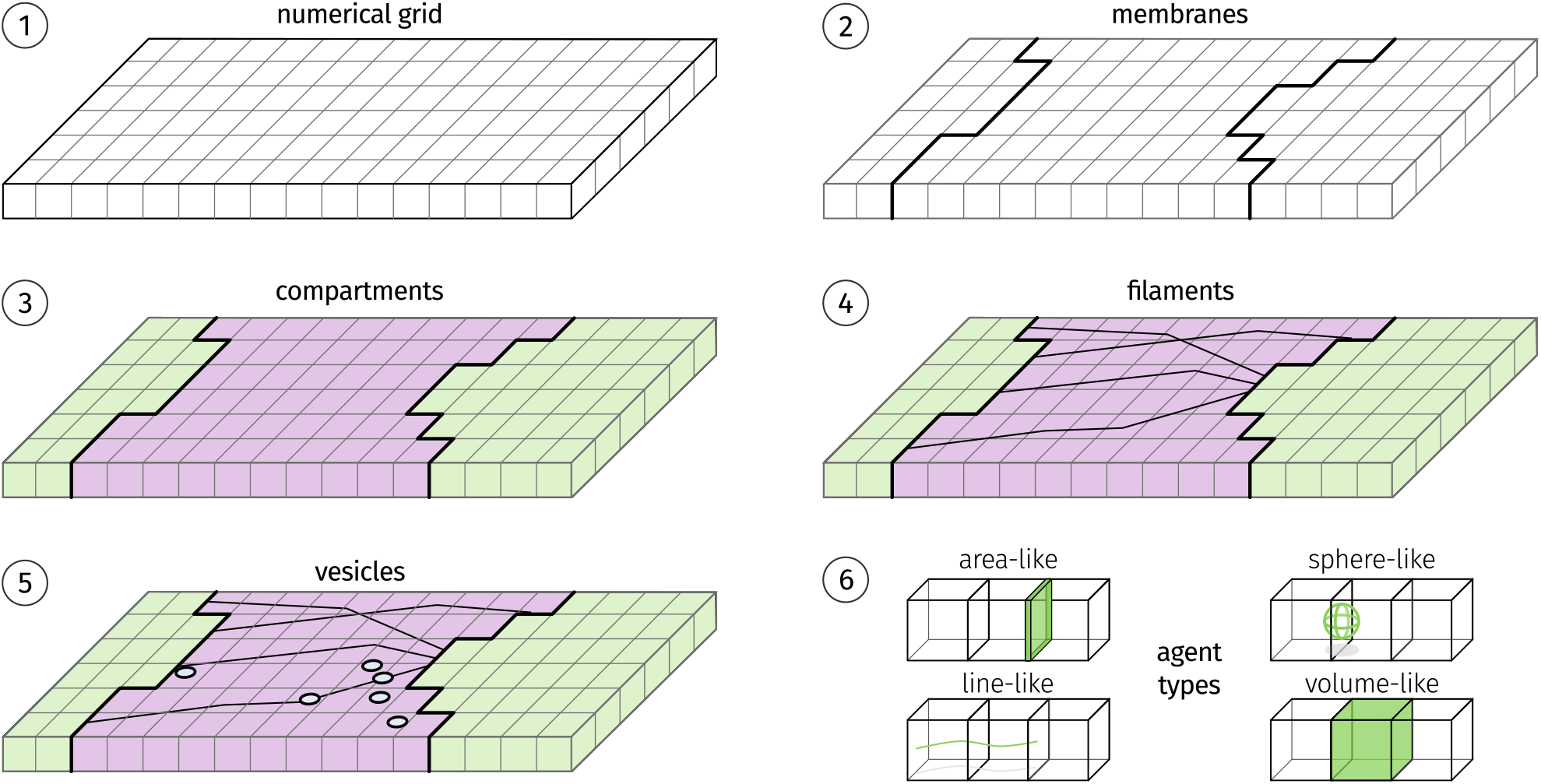
Components of the simulation. The simulation space (**1**) is tiled into a regular grid, used for numerical calculation and spatial indexing. The membrane agents (**2**) determine the compartments (**3**) of the simulation. Each cell of the grid is assigned a concentration of chemical entities. Filaments (**4**) and vesicles (**5**) are placed in the simulation. All agents are implementations of abstract agent types (**6**).

Macroscopic entities are modeled as agents that are able to interact with each other and with the reaction spaces (see Figure 7). The interactions are defined by a set of conditions that have to be met before associated actions are performed. Vesicles are sphere-like agents able to move across the simulation space. A vesicle posses a state, a position, a radius and two compartments.

The compartments represent the internal cargo and the membrane surface. The state of a vesicle determines how it is processed by modules. Since vesicles move, they have to change their neighboring compartments dynamically during simulation. This entails that each vesicle is referenced to up to four additional compartments. If a vesicle is on the border between a number of grid cells the area of the intruding membrane section is calculated and used to scale the exposed concentrations of chemical entities. The aforementioned membranes are area-like agents, made up of membrane segments. Each segment is represented by a line between two membrane separated grid points. This allows for the calculation of the membrane area per compartment and is used to scale the amount of reactants available to reactions. Cytoskeletal filaments are line-like agents for the directed transport of vesicles. Line-like agents are composed of multiple segments and possess a positive and a negative end to indicate a direction. This directionality is used for the transport of vesicles that are attached via chemical entities that are considered molecular motors. Volume-like agents are used impose specific behavior to a subsection of the simulation space. For example, they are used to define the cell cortex, where vesicles are tethered to the membrane.

### Module based update system

The simulation integrates changes over time that are calculated by independent components called modules. A module can be either concentration-based, displacement-based or qualitative. Concentration based modules determine changes in concentrations of chemical entities resulting from reactions and transport processes. Displacement based modules are used to move agents inside of the cell. Qualitative modules can implement rules and algorithms, for example deciding whether a vesicle should be attached to a filament or when and how vesicles fuse with membranes.

The reactions between chemical entities are defined using a rule-based system. The definition of reactions uses a combination of modification operations: binding, release, addition, or removal. The modification operations basically describe which parts of a chemical entity are added or removed by a reaction. Additionally, criteria can be defined that narrow the amount and kind of chemical entities that are able to react (see Fig. 8). A network generation algorithm determines all possible reactions that can occur with the defined rules and return them to the user for possible refinement. The result of the network generation is a set of ordinary differential equations. Each equation represents an elementary chemical reaction that follows the law of mass action. The behavior of each compartment is determined by these equations that define concentration change of each chemical entity. Additionally, transport processes use the concentrations of multiple adjacent compartments to determine a change in concentration^92^.

**Figure 8.**
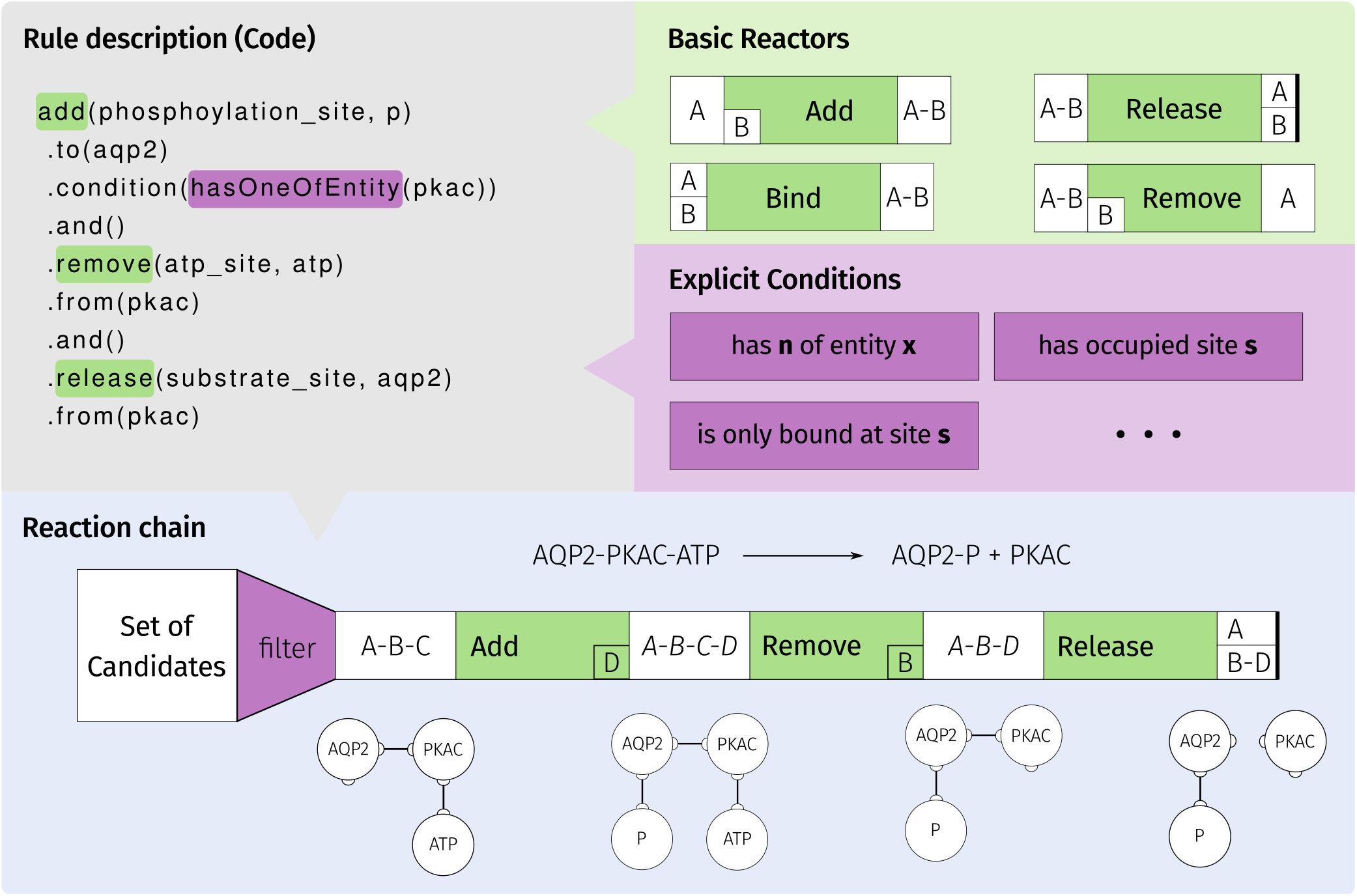
Definition of reaction rules. Reactions can be specified by a description of the reaction process. Basic reactors are concatenated to chains that are processed during the reaction network generation. Additionally, conditions allow for further specification of the reaction process. The network generation process results in all reactions that can occur in the system.

The sum of changes for each entity and compartment is determined by the sum of reactions that affect a chemical entity. A local numerical error is calculated on a reaction basis. The error is used to decrease or increase the time step and therefore determines the trade-off between accuracy and speed of the simulation. The evaluation on reaction level allows for the determination of reactions with high numerical error, which in turn allows for their individual optimization until a stable time step is found. A detailed description of the error calculation can be found in the Supplementary Methods. After all local errors are acceptable, a total numerical error is calculated that evaluates the change on chemical entity level.

Displacement based modules are implemented using a similar approach. The maximal displacement should be small, since abrupt compartment changes for vesicles can lead to instability in the numerical computations. Therefore, the maximal displacement is set to a fraction of the numerical grid step width. A local displacement per module and a total displacement per vesicle is calculated. If the displacement is too large, the time step is decreased, if it is comparably small the time step can be increased. Quantitative modules are able to implement functions that evaluate the error of the function and are equally able to request time step decrease and increase from the simulation.

The modular system in combination with individual error management allows for an extensible framework that enables the simulation of multiscaled cellular systems coupling macroscopic and microscopic environments.

## Supplementary information

### Supplementary Methods

Contains additional and detailed descriptions of the modeling and simulation approach devised in this publication.

### Supplementary Information 1

Detailed model description. Contains background, details, parameters, and corresponding literature used for the derived models.

### Supplementary Information 2

Estimation of AQP2 concentrations in membrane and vesicles during the signalling progress.

**Supplementary Figure 1** Overview of the processes and reactions modeled in SBGN notation. Micro-scale and reaction based processes and entities are colored blue, whereas macroscopic and agent based processes are colored green.

## Author contributions statement

C.L. devised the models, wrote the simulation software, analyzed the resulting data, and prepared the figures. C.L. wrote the manuscript. D.L. and M.S supervised the project. All authors read and approved the manuscript.

## Additional information

The authors declare no conflict of interest.

## Acknowledgements

We would like to thank Floian Kaiser, Sebastian Bittrich, as well as Alexander Eisold for valuable discussions and proofreading the manuscript. This work is supported by the European Social Fund (ESF).

